# SH3Ps recruit auxilin-like vesicle uncoating factors into clathrin-mediated endocytosis

**DOI:** 10.1101/2022.01.07.475403

**Authors:** Maciek Adamowski, Ivana Matijević, Madhumitha Narasimhan, Jiří Friml

## Abstract

Clathrin-mediated endocytosis (CME) is an essential process of cellular cargo uptake operating in all eukaryotes. In animal and yeast, CME involves BAR-SH3 domain proteins, endophilins and amphiphysins, which function at the conclusion of CME to recruit factors for vesicle scission and uncoating. *Arabidopsis thaliana* contains BAR-SH3 domain proteins SH3P1-3, but their role is poorly understood. We identify SH3P1-3 as functional homologues of endophilin/amphiphysin. SH3P1-3 bind to discrete foci at the plasma membrane (PM), and colocalization indicates late recruitment of SH3P2 to a subset of clathrin-coated pits. PM recruitment pattern of SH3P2 is nearly identical to its interactor, a putative vesicle uncoating factor AUXILIN-LIKE1, and SH3P1-3 are required for most of AUXILIN-LIKE1 PM binding. This indicates a plant-specific modification of CME, where BAR-SH3 proteins recruit auxilin-like uncoating factors, rather than the uncoating phosphatases synaptojanins. Furthermore, we identify an unexpected redundancy between SH3P1-3 and a plant-specific endocytic adaptor, TPLATE complex, showing a contribution of SH3P1-3 to gross CME.

## Introduction

Endocytosis is a process at the plasma membrane (PM) leading to the internalization of surface proteins and other cargo. In plants, as in other eukaryotes, the most common mode of endocytosis is a vesicular transport process involving the vesicle coat protein clathrin (Dhonukshe et al., 2007; Narasimhan et al., 2020). Clathrin-mediated endocytosis (CME) has been studied in detail in animals and yeast (Kaksonen and Roux, 2018), describing it as a process of high complexity involving, beyond clathrin, a large array of other protein factors (Merrifield and Kaksonen, 2014). These perform functions that include binding clathrin with the membrane, force generation for membrane bending, and selective loading of cargo at earlier stages. At late stages of CME, specialized factors mediate the separation of the completed vesicle from the PM and uncoating, i.e., removal of the protein coat to release the vesicle for fusion with an endosomal compartment.

In non-plants, the formation of clathrin-coated vesicles (CCVs), especially the late steps leading to scission and uncoating, involves the action of BAR (Bin, amphiphysin and Rvs) domain proteins (Nishimura et al., 2018; Simunovic et al., 2019). BAR domains are dimers forming structures of a crescent shape, which typically bind membranes through concave surfaces displaying positive charges. As such, BAR domains act as curvature sensors, binding, for instance, to narrow necks of nearly-formed CCVs, and as curvature generators, where their binding and polymerization leads to further membrane bending. CME involves a suite of BAR domain proteins with distinct recruitment characteristics and composition beside the BAR domain (Taylor et al., 2011; Carman and Dominguez, 2018). Notable examples are amphiphysins and endophilins, which possess an SH3 domain, an interaction module with affinity for Proline-rich sequences on protein targets (Kurochkina and Guha, 2013). The SH3 domains of amphiphysins and endophilins interact with dynamins, mechanoproteins acting in vesicle scission, and with synaptojanins, phospholipid phosphatases that promote uncoating by modifying vesicle membrane lipid composition (Ringstad et al., 1997; Grabs et al., 1997; Micheva et al., 1997). Thus, thanks to curvature-sensing properties of BAR domains and the interactions of SH3 domains, BAR-SH3 proteins act as binding intermediates enabling a temporally regulated activity of factors needed for the final steps of CME (Shupliakov et al., 1997; Verstreken et al., 2003; Schuske et al., 2003; Milosevic et al., 2011).

The characterization of CME in *Arabidopsis thaliana*, including the steps leading to scission and uncoating, is less advanced (Reynolds et al., 2018). *A. thaliana* contains three proteins with a BAR-SH3 domain composition similar to non-plant endophilin/amphiphysin, SH3P1 to SH3P3 (SH3 DOMAIN-CONTAINING PROTEIN1-3). In support of a conserved function, expression of SH3P1-3 partially complements the yeast amphiphysin mutant *rvs167* (Lam et al., 2001). Interactions between SH3P1-3 and plant dynamins were detected (Lam et al., 2002; Ahn et al., 2017; Baquero Forero and Cvrcková, 2019). With regard to uncoating factors, *A. thaliana* lacks evident homologues of synaptojanin, but interestingly, SH3Ps interact with some of the putative *A. thaliana* homologues of auxilin, a class of uncoating factors that act by the recruitment of Hsc70 (heat shock cognate 70), causing relaxation of the clathrin cage in mammals (Sousa and Lafer, 2015). The homologous AUXILIN-LIKE1 and AUXILIN-LIKE2 possess Pro-rich domains and interact with SH3Ps, as detected by yeast two-hybrid, co-immunoprecipitation, bimolecular fluorescence complementation, and common isolation in Tandem Affinity Purification with CLATHRIN LIGHT CHAIN (CLC) as bait (Lam et al., 2001; Adamowski et al., 2018). These interactions may represent a novel, plant-specific modification of CME, where BAR-SH3 proteins engage with auxilin-like uncoating factors, rather than with synaptojanins, to promote CCV uncoating (Lam et al., 2001), but functional consequences of this model were not explored *in planta*. Overall, a comprehensive evaluation of the role of SH3P1-3 in CME is lacking. That said, functions such as trafficking of ubiquitinated cargoes, autophagosome formation, and cell plate formation, were assigned to SH3P2 (Zhuang et al., 2013, Ahn et al., 2017, Nagel et al., 2017).

Here, with the use of loss-of-function and advanced live imaging approaches, we characterize the role of SH3P1-3 in CME, and explore their connections with the putative uncoating factors AUXILIN-LIKE1/2. Our results demonstrate a role of SH3P1-3 as homologues of endophilin/amphiphysin, where they engage in a novel, plant-specific association with auxilin-like uncoating factors. This study contributes to the on-going characterization of the unique properties of CME, which evolved in the plant lineage.

## Results

### Isolation and complementation of *sh3p123* mutants

Literature reports variably on the effects of SH3P1-3 deficiency. Multiple *sh3p* mutant combinations, with T-DNA alleles that are likely not full knock-outs, did not exhibit obvious phenotypic defects (Ahn et al., 2017; Nagel et al., 2017). In turn, an RNA interference (RNAi) line silencing *SH3P2* exhibited an arrest of seedling development (Zhuang et al., 2013), indicating an essential and non-redundant function of SH3P2, but this was not reproduced with a similar artificial microRNA (amiRNA) line (Nagel et al., 2017).

To clarify the function of SH3P1-3, we aimed to isolate *sh3p123* triple mutants using CRISPR/Cas9-generated mutations in the exons of *SH3P1* and *SH3P2*, and an exon T-DNA insertion in *SH3P3*, all of which disrupt coding sequences, producing null alleles. We obtained two independent triple homozygous allele combinations, named *sh3p123*^*9C*^ and *sh3p123*^*10G*^. The triple knock-outs exhibited reduced growth as seedlings (Figure 1A) as well as adults (Figure 1B) and had limited fertility. These mutant phenotypes were absent from *sh3p12* and *sh3p3* (Suppl. Figure 1A and 1B), indicating a functional redundancy of the three isoforms. We complemented the triple mutant by transforming *sh3p123*^*10G*^ with *35S*_*pro*_*:SH3P1-GFP, 35S*_*pro*_*:SH3P2-GFP*, and *35S*_*pro*_*:SH3P3-GFP*. Expression of any of the three fusion proteins successfully complemented the mutant to various degrees in individual transgenic lines (Figure 1C). This confirms that the three isoforms possess a redundant function, and also shows functionality of C-terminal fluorescent protein fusions.

**Figure 1.**
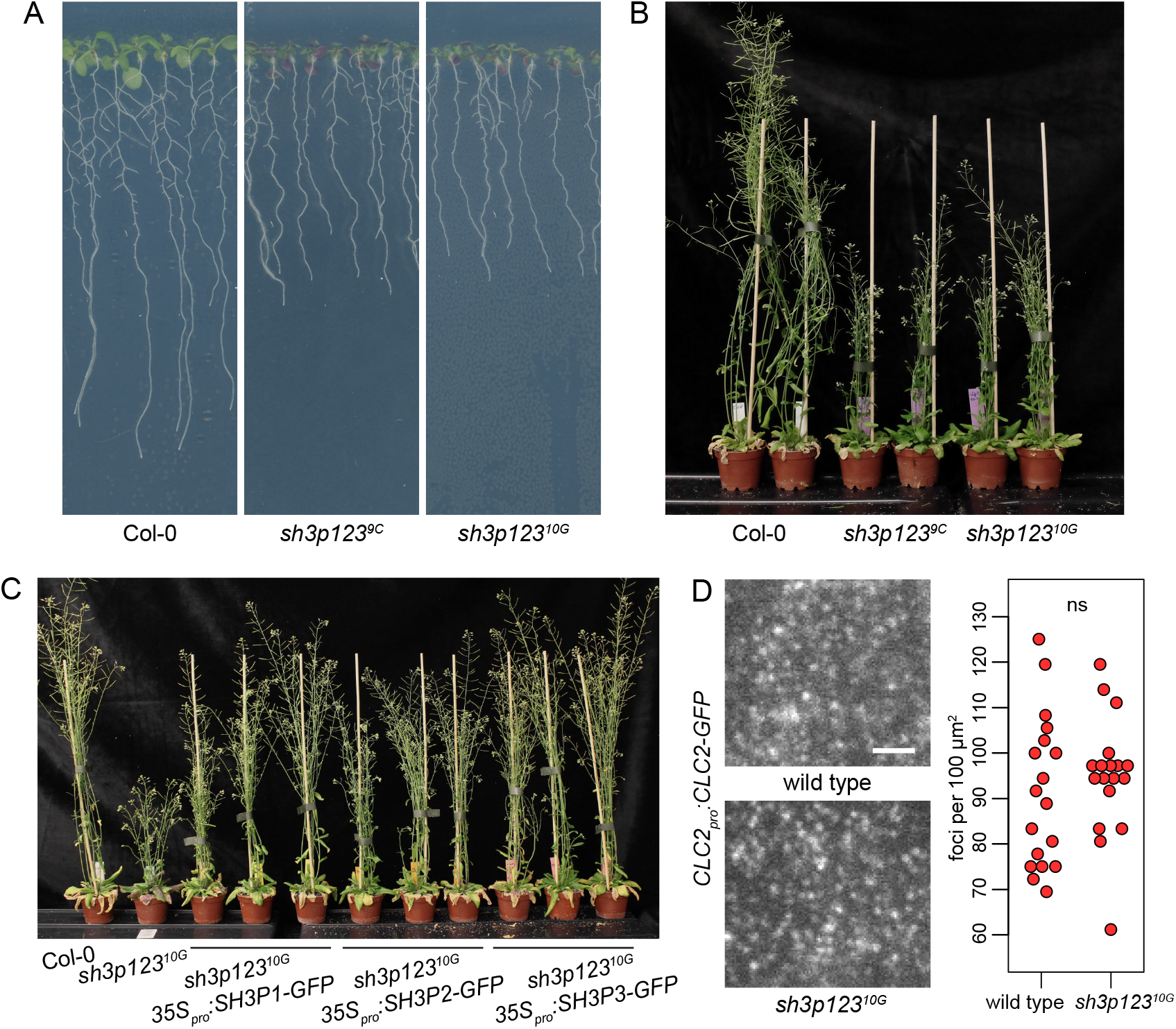
sh3p123 loss of function mutants (A, B) Seedling (A) and adult (B) development of sh3p123 knock-out mutants. Two independently obtained allele combinations are shown. sh3p123 mutants are characterized by a reduced growth of seedlings and adults. (C) Complementation of sh3p12310G with 35Spro:SH3P1-GFP, 35Spro:SH3P2-GFP, and 35Spro:SH3P3-GFP. Single plants from three independent transgenic lines carrying each constructs are shown. All three constructs complement the mutant phenotype with varying degrees in individual lines. (D) TIRF images of CLC2-GFP in the epidermis of early elongation zone of seedling roots of sh3p12310G. Graph shows quantifications of CLC2-GFP-positive foci density, each data point representing one root. Wild type: 91.4 ± 16.7 foci per 100 μm2 (mean ± s.d.), n=18; sh3p12310G: 94.9 ± 13.1 foci per 100 μm2, n=18. Values were compared using a t test, ns – not significant.

To test the requirement of SH3P1-3 for CME, we crossed *sh3p123*^*10G*^ with *CLC2*_*pro*_*:CLC2-GFP UBQ10*_*pro*_*:mCherry-AUXILIN-LIKE1* reporter line. We evaluated CME by Total Internal Reflection Fluorescence (TIRF) of CLC2-GFP, a clathrin coat marker. Measured in the epidermis of the early elongation zones of seedling roots, the densities of clathrin-coated pits (CCPs) at the PMs were equal in the wild type and the mutant (Figure 1D), indicative of grossly normal CME in the absence of SH3P1-3.

Taken together, knock-out mutants and their complementation indicate that SH3P1-3 possess a redundant function supporting growth and development, but contribution of SH3P1-3 to gross CME was not detected.

### Localization of SH3P1-3 at the PM and recruitment to CCPs

Subcellular localization data places SH3P2-GFP at the PM and at cell plates, on endomembrane structures, and on autophagosomes (Zhuang et al., 2013; Ahn et al., 2017; Nagel et al., 2017; Adamowski et al., 2018). The association of SH3P2 with the PM is mediated by the BAR domain (Nagel et al., 2017). We evaluated the localization of all three SH3Ps using functional SH3P-GFP fusions expressed in complemented *sh3p123*^*10G*^. Observed with confocal laser scanning microscopy (CLSM) in seedling root apical meristem (RAM), the major site of SH3P1-GFP, SH3P2-GFP, and SH3P3-GFP localization was the PM (Figure 2A). All three fluorescent protein fusions localized to cell plates during cell division (Suppl. Figure 1C). TIRF imaging of the PM localization patterns of all three isoforms showed relatively sparse, clear fluorescent foci of signal which persisted at the PM for varied, but generally short, amounts of time (Figure 2B).

**Figure 2.**
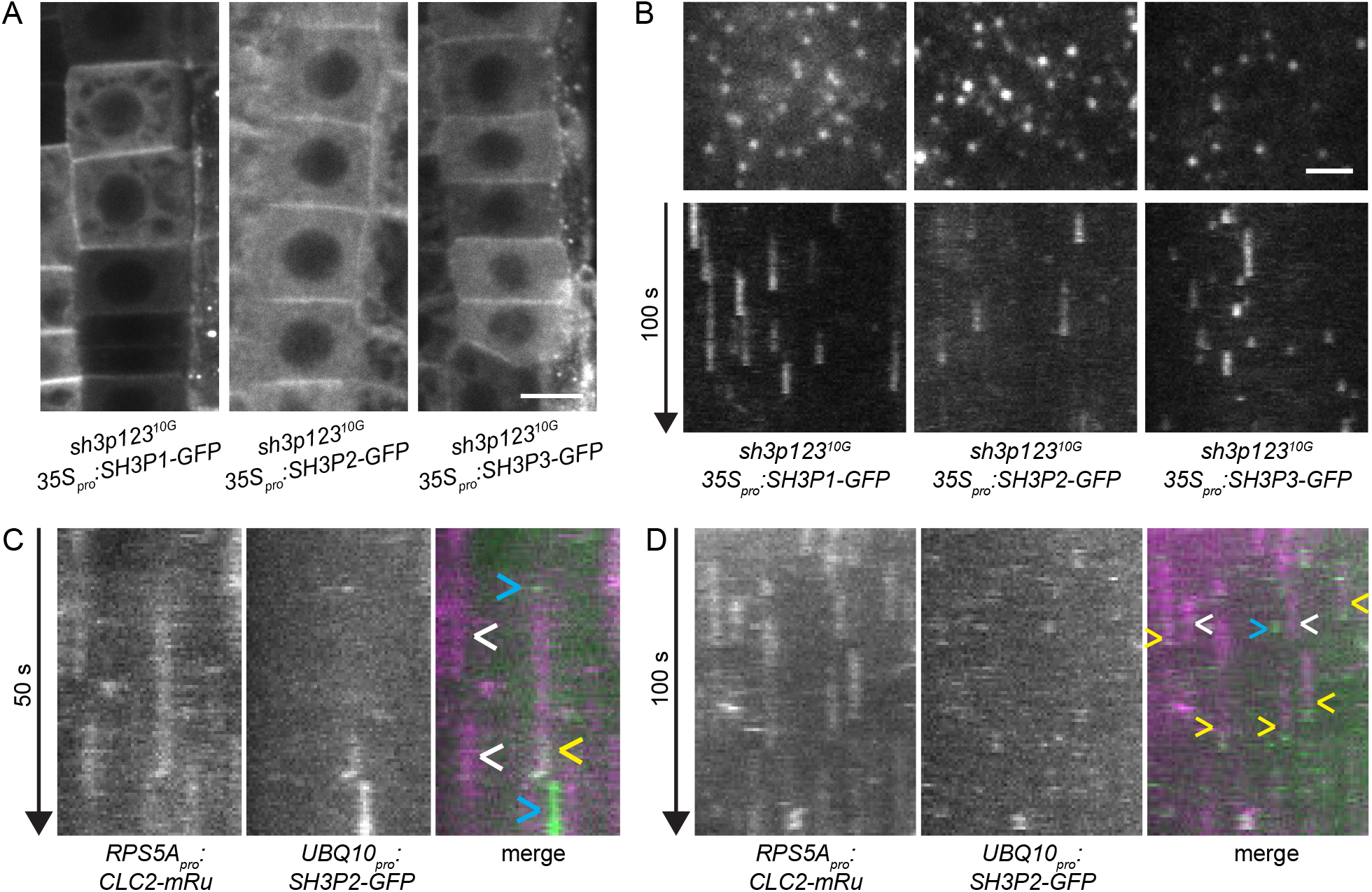
PM localization of SH3P1-3 and recruitment to CCPs (A) CLSM images of SH3P1-GFP, SH3P2-GFP, and SH3P3-GFP in RAM epidermis of complemented sh3p12310G mutants. All fluorescent protein fusions are enriched at the PMs. Scale bar – 10 μm. (B) Single TIRF frames (top) and kymographs (bottom) of TIRF time lapses of SH3P1-GFP, SH3P2-GFP, and SH3P3-GFP in the epidermis of early elongation zone of seedling roots in complemented sh3p12310G mutants. All fluorescent protein fusions localize to the PM as distinct foci, which persist for varied amounts of time. Scale bar – 2 μm. (C, D) Kymographs of TIRF time lapse colocalization between CLC2-mRuby3 and SH3P2-GFP in the epidermis of root early elongation zone (C) and of etiolated hypocotyls (D). SH3P2-GFP is recruited at the end of some instances of CCV formation (yellow arrowheads). In (C), the completed CCV with SH3P2-GFP can be seen departing from the PM to the left in the plane of section, before the signal disappears presumably due to vesicle uncoating. Many captured CCPs do not recruit SH3P2-GFP (white arrowheads). SH3P2-GFP is also recruited at sites not containing clathrin (blue arrowheads).

To assess if SH3P1-3 localize at CCPs, we generated a double marker line *UBQ10*_*pro*_*:SH3P2-GFP RPS5A*_*pro*_*:CLC2-mRuby* and conducted TIRF time lapse imaging in the early elongation zone of the root and in etiolated hypocotyls. In both tissues, we found SH3P2-GFP traces associated with a subset of CCPs marked by CLC2-mRuby, where SH3P2-GFP was recruited at the final stage of CCP formation that leads to the detachment of the CCV from the PM (Figure 2C and 2D, yellow arrowheads). This colocalization pattern indicates SH3P1-3 function at late stages of CCP formation leading to scission and uncoating, similarly to endophilins and amphiphysins (Taylor et al., 2011). Beside these colocalizations, many CCP formations did not involve SH3P2-GFP (Figure 2C and 2D, white arrowheads), and instances of SH3P2-GFP localization at sites not containing clathrin were seen (Figure 2C and 2D, blue arrowheads).

In summary, SH3P1-3 dynamically localize to distinct spots at the PM, and imaging of SH3P2-GFP indicates their late recruitment to a subpopulation of CCPs, suggestive of a potential function in late stages of CME, similar to endophilin and amphiphysin in non-plant systems.

### SH3P1-3 associate with AUXILIN-LIKE1/2 at the PM

In non-plant systems, endophilin and amphiphysin interact at CCPs with Pro-rich domain-containing partners, synaptojanin and dynamin (Ringstad et al., 1997; Grabs et al., 1997; Micheva et al., 1997). Protein-protein interactions support the notion that SH3P1-3 engage with the putative uncoating factors AUXILIN-LIKE1/2 (Lam et al., 2001; Adamowski et al., 2018), possibly to recruit them to CCPs. In TIRF imaging studies, AUXILIN-LIKE1 was observed binding to a subset of CCPs at late stages of formation (Adamowski et al., 2018). Additionally, AUXILIN-LIKE1 associated with the PM at sites not containing clathrin. Overall, the localization pattern and dynamics of AUXILIN-LIKE1 at the PM were very similar to those of SH3P2.

To test if SH3P2 and AUXILIN-LIKE1 localize at the PM together, indicative of physical and functional associations between SH3P1-3 and AUXILIN-LIKE1/2, we generated a double marker line *UBQ10*_*pro*_*:SH3P2-GFP UBQ10*_*pro*_*:mCherry-AUXILIN-LIKE1*. In TIRF microscopy, the two fluorescent markers revealed a striking degree of co-localization, where in most instances, they arrived and left the PM simultaneously at the same locations (Figure 3A, yellow arrowheads). Some instances of SH3P2-GFP recruitment without associated mCherry-AUXILIN-LIKE1 could be seen (Figure 3A, blue arrowheads). The high degree of co-localization strongly indicates that SH3P1-3 interact with AUXILIN-LIKE1/2 at the PM. Given that a fraction of both these proteins is found at sites distinct from clathrin (Figure 2C and 2D; Adamowski et al., 2018), many of the interactions may not take place at CCPs, but it is likely that common recruitment and interactions between SH3P1-3 and AUXILIN-LIKE1/2 occur at CCPs as well.

**Figure 3.**
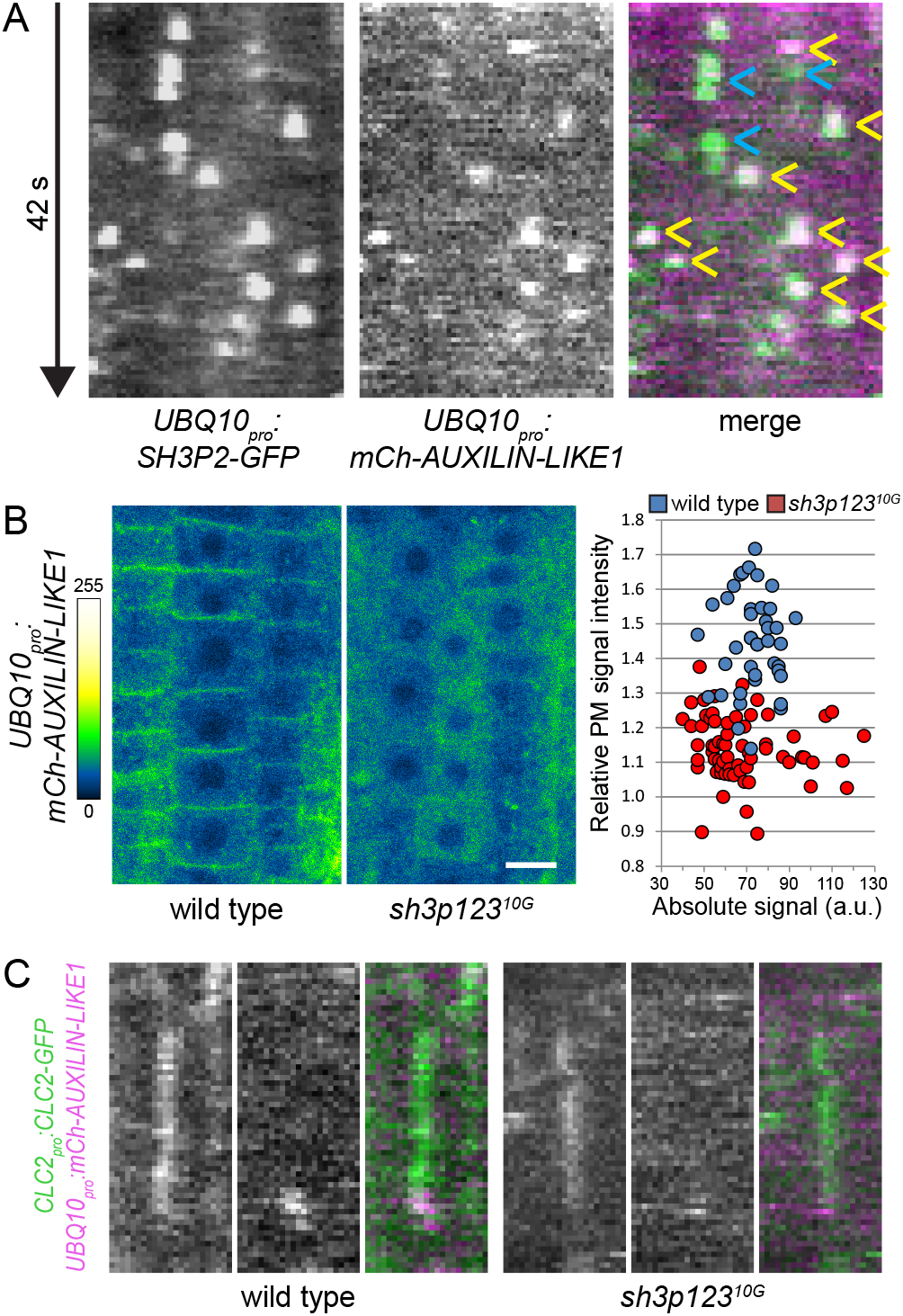
SH3P1-3 recruit AUXILIN-LIKE1/2 to the PM (A) Kymographs of TIRF time lapse colocalization between SH3P2-GFP and mCherry-AUXILIN-LIKE1 in the epidermis of etiolated hypocotyls. The two fluorescent protein fusions exhibit a high degree of colocalization at structures of typically relatively brief life times (yellow arrowheads). SH3P2-GFP is also recruited at sites not containing mCherry-AUXILIN-LIKE1 (blue arrowheads). (B) CLSM images of mCherry-AUXILIN-LIKE1 in wild type and sh3p123 seedling RAM epidermis. The localization of mCherry-AUXILIN-LIKE1 to the PM is almost completely lost in the mutant. Scale bar – 10 μm. Graph shows relative PM signal intensities in individual seedlings and lack of their correlation with absolute signal levels. Relative PM signal intensities: wild type 1.45 ± 0.14 (mean ± s.d.), n=41, sh3p123 1.14 ± 0.09, n=65. Values were compares using a t test, P<0.0001. (C) Kymographs of TIRF time lapse colocalization between CLC2-GFP and mCherry-AUXILIN-LIKE1 in the epidermis of etiolated hypocotyls of wild type and sh3p123. Events of recruitment of mCherry-AUXILIN-LIKE1 at the end of CCV formations can be observed in both genotypes.

The most likely scenario linking SH3P1-3 and AUXILIN-LIKE1/2 at the PM is a direct membrane binding of SH3P1-3 through BAR domains, and a recruitment of AUXILIN-LIKE1/2 by Pro-rich-SH3 domain interaction. To test if SH3P1-3 are required for AUXILIN-LIKE1/2 localization at the PM, we imaged mCherry-AUXILIN-LIKE1 in *sh3p123*^*10G*^ background. When assessed with CLSM in seedling RAM epidermis, the PM-associated signals of mCherry-AUXILIN-LIKE1 were virtually lost in the mutant (Figure 3B), and a major portion of the protein was present in the cytosol. The relative mCherry-AUXILIN-LIKE1 signal intensity at the PM was not correlated with absolute expression levels in individual seedlings (Figure 3B). This experiment indicates that the PM recruitment of AUXILIN-LIKE1/2 is mediated by SH3P1-3, consistent with their physical interactions and common spatio-temporal occurrence at the PM. We analysed this in further detail by co-localizing mCherry-AUXILIN-LIKE1 with CLC2-GFP in *sh3p123*^*10G*^ by TIRF microscopy, to verify if recruitment of AUXILIN-LIKE1 at the conclusion of CCP formation occurs in the absence of SH3P1-3. While a quantitative comparison between genotypes was not feasible due to the rarity of reliable detection of these events (Adamowski et al., 2018), some such co-localizations could be decisively observed both in the mutant, and in the wild type (Figure 3C). Putting all the observations together, we conclude that SH3P1-3 clearly possess a function in the PM recruitment of AUXILIN-LIKE1/2, but an additional mechanism of their recruitment to CCPs likely operates.

### Distinct responses of SH3P2 to CME inhibition by clathrin silencing and AUXILIN-LIKE1 overexpression

Overexpression of AUXILIN-LIKE1/2, or silencing *CLATHRIN HEAVY CHAIN (CHC)* with amiRNA, leads to an inhibition of CME (Adamowski et al., 2018; Adamowski and Friml, 2021). In both cases, interference with CME manifests similarly in terms of the endocytic machinery. In each case, the recruitment of CLC2-GFP to CCPs is diminished. Subunits of early acting TPLATE complex and AP-2 (ADAPTOR PROTEIN 2) (Gadeyne et al., 2014) remain at the PM and exhibit elevated binding, indicating that inhibition of CME occurred later, at the stage of clathrin recruitment. In both cases, too, the dynamin DRP1C-GFP showed a partial, variable signal decrease (Adamowski et al., 2018; Adamowski and Friml, 2021).

We tested how SH3P2-GFP responds to inhibition of CME in these two lines. Silencing of *CHC* in *XVE»amiCHCa* lead to a loss of SH3P2-GFP from the PM in almost all seedlings analysed by CLSM (Figure 4A). Similarly, TIRF in early elongation zone of the root showed loss of most of the punctate PM signals, and SH3P2-GFP was found preferentially in the cytosol (Figure 4C, middle). This supports a function of SH3P1-3 as late-acting components of CCPs. Given that SH3P2-GFP is often present at sites not containing clathrin, this recruitment may have been abolished in *XVE»amiCHCa* as well, for unknown reasons. Interestingly, the response of SH3P2-GFP to AUXILIN-LIKE1 overexpression was different: CLSM revealed that SH3P2-GFP not only remained at the PM, but the relative PM binding levels were higher than in control conditions (Figure 4B). Consistently, TIRF showed abundant, often enlarged foci of SH3P2-GFP signal (Figure 4C, right).

**Figure 4.**
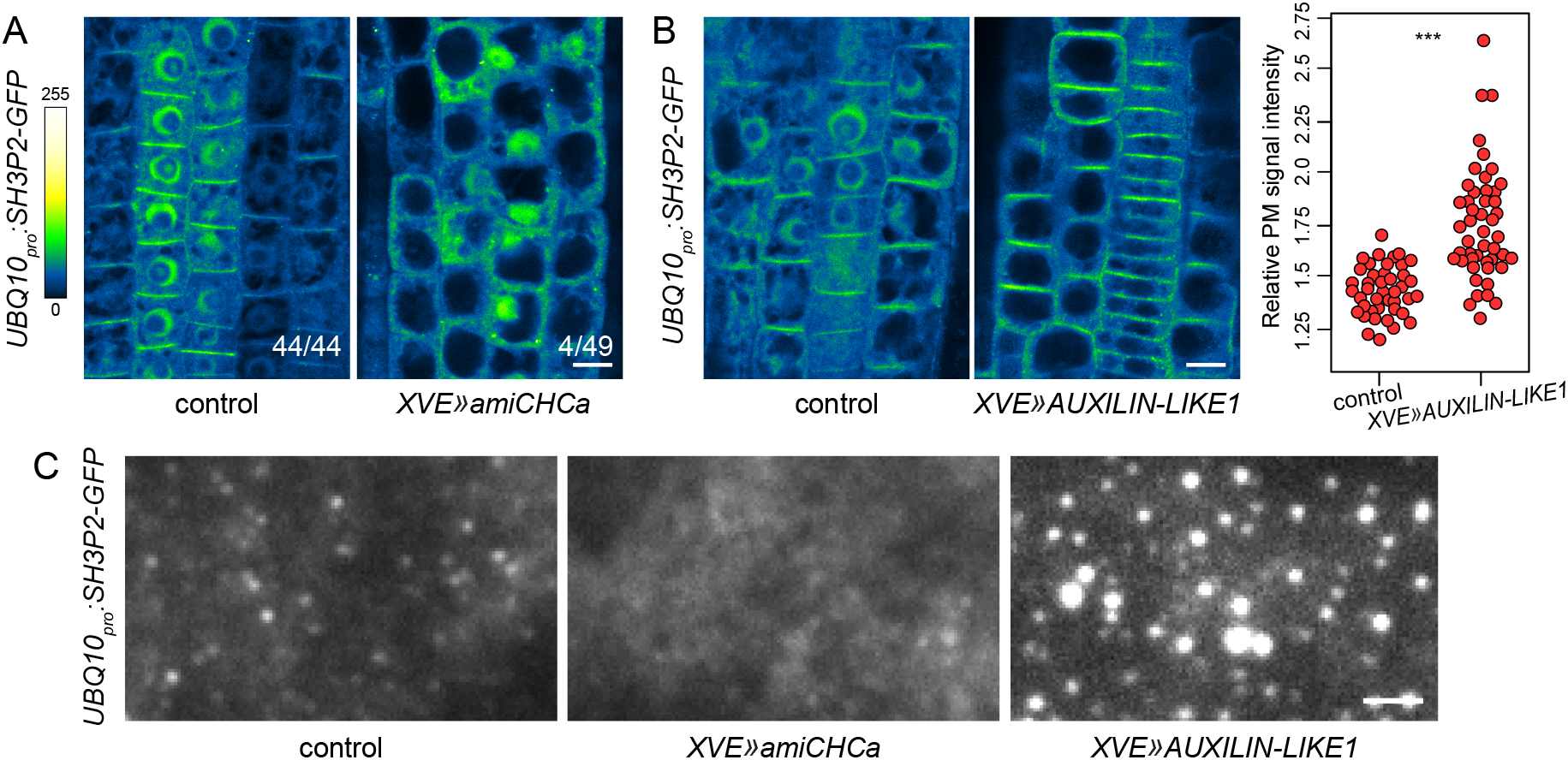
A distinct reaction of SH3P2-GFP to CME inhibition by AUXILIN-LIKE1 overexpression compared with clathrin silencing (A)CLSM images of SH3P2-GFP in seedling RAM epidermis of XVE»amiCHCa induced for approx. 48 h. Silencing of CHC leads to a loss of SH3P2-GFP signals from the PM. Numbers indicate ratio of RAMs with PM signals of SH3P2-GFP. Scale bar – 10 μm. (B) CLSM images of SH3P2-GFP in seedling RAM epidermis of XVE»AUXILIN-LIKE1 induced for approx. 24 h. Overexpression of AUXILIN-LIKE1 leads to variable increases in the PM signal relative to total RAM signals. Scale bar – 10 μm. Graph shows a quantification of relative PM signal intensities, each data point representing one root. Control: 1.43 ± 0.12 (mean ± s.d.), n=42; XVE»AUXILIN-LIKE1: 1.75 ± 0.27, n=48. Values were compared using a t test, P<0.0001. (C) TIRF images of SH3P2-GFP in seedling root epidermis of XVE»amiCHCa and XVE»AUXILIN-LIKE1 induced for approx. 48 h and 24 h, respectively. Following CHC silencing, SH3P2-GFP is observed as cytosolic background with only rare clear foci of signal. Overexpression of AUXILIN-LIKE1 causes an accumulation of SH3P2-GFP signal in abundant foci which are often enlarged. Scale bar – 2 μm.

The distinct response of SH3P2-GFP to AUXILIN-LIKE1 overexpression compared with *CHC* silencing, characterized by elevated PM binding, supports unique functional connections of SH3P1-3 and AUXILIN-LIKE1/2, and additionally indicates that inhibition of CME by AUXILIN-LIKE1/2 overexpression proceeds by a specific mechanism.

### Functional relationship of SH3P1-3 and the TPLATE complex

Our analysis of SH3P1-3 indicates their activity at CCPs, and functional connections with the putative uncoating factors AUXILIN-LIKE1/2. Yet, the deleterious consequences of *sh3p123* loss-of-function are relatively mild, and not associated with a detectable deficiency in CME as a whole. We speculated about a possible functional redundancy between SH3P1-3 and another, unknown component. The only other protein with a predicted SH3 domain in *A. thaliana* (our BLAST search), which, incidentally, also functions in CME, is the TPLATE complex subunit TASH3 (TPLATE ASSOCIATED SH3 DOMAIN CONTAINING PROTEIN) (Gadeyne et al., 2014). The TPLATE complex, belonging to the TSET complex class, is structurally homologous to the tetrameric adaptor protein complexes AP-1 through AP-5, some of which act as clathrin adaptors (Sanger et al., 2019). TASH3 is in this context a homologue of the γαδεζ large subunit of APs, but uniquely, contains an SH3 domain at the C-terminus, absent not only in AP complexes, but also in non-plant TSET (Hirst et al., 2014).

To assess if the SH3 domain of TASH3 functionally overlaps with those of SH3P1-3, we isolated *tash3* mutants, and introduced them into *sh3p123*. Mutants of the TPLATE complex subunits exhibit male gametophytic lethality, precluding a straightforward loss-of-function analysis (Gadeyne et al., 2014). Indeed, we could not recover homozygotes of *tash3-1* and *tash3-2* lines harbouring T-DNA insertions in early exons of *TASH3* (Figure 5A), presumably due to lethality of mutant male gametes. Instead, given that the SH3 domain of TASH3 is C-terminal, we attempted to isolate mutants where only the sequence coding for the SH3 domain is disrupted. This may lead to the expression of a truncated TASH3 protein containing all parts of structure homologous to subunits of AP complexes, possibly still retaining partial functionality. We obtained mutants of this kind by two approaches. First, we isolated *tash3-3*, an exon T-DNA insertion shortly before the sequence encoding the SH3 domain (Figure 5A). Conceptual translation based on re-sequencing of the T-DNA insertion border predicted a TASH3 protein terminated before the SH3 domain (Suppl. Figure 2A). Additionally, we targeted *TASH3* by CRISPR/Cas9, generating a *tash3-c* allele with a single nucleotide insertion early in the SH3 domain-encoding sequence, leading to a stop codon within this domain in the predicted protein (Figure 5A and Suppl. Figure 2A).

**Figure 5.**
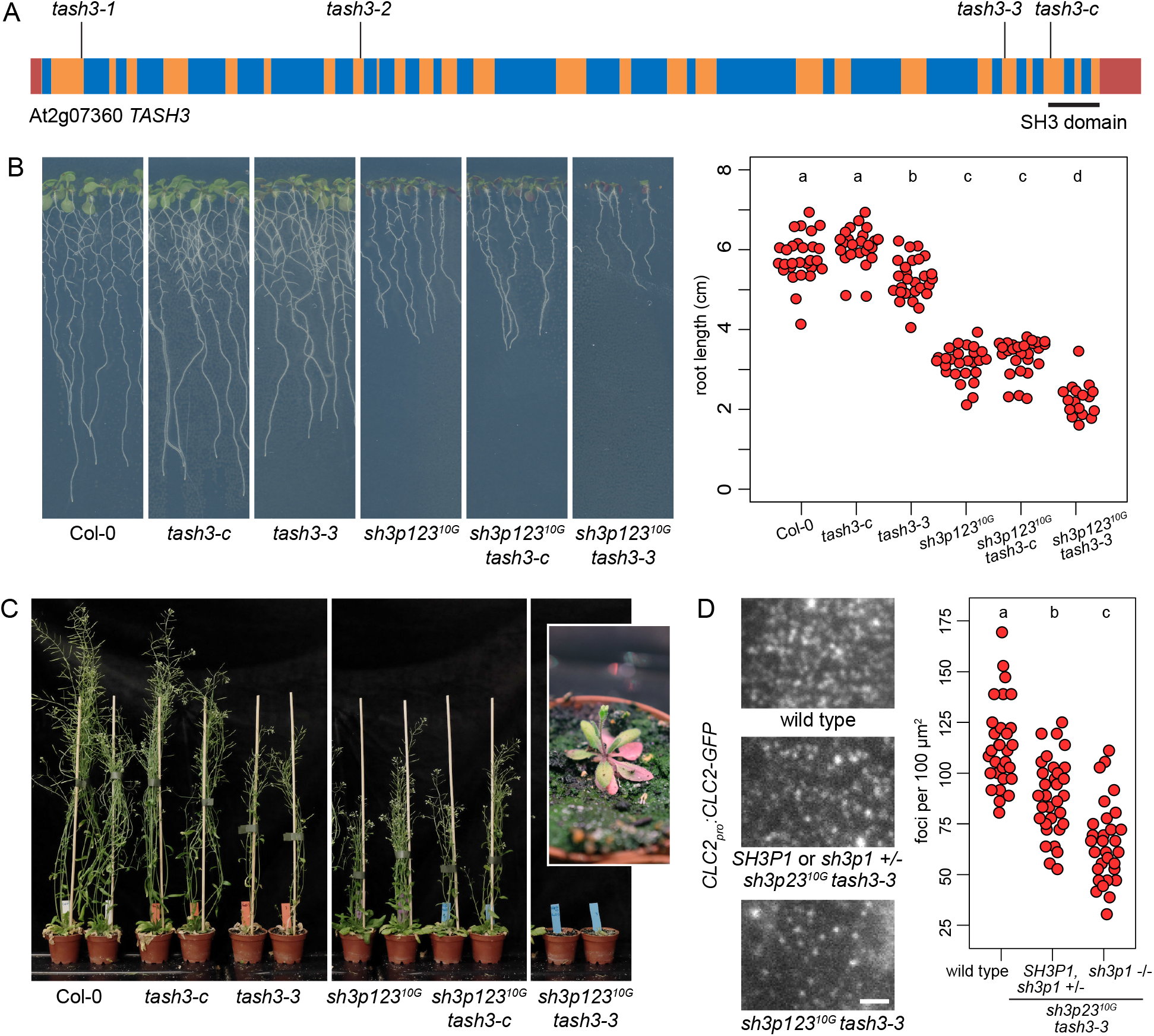
Functional connections between SH3P1-3 and TASH3 (A) TASH3 gene structure and mutant alleles. Exons are represented in orange, introns in blue, and UTRs in red. Sequence encoding the C-terminal SH3 domain is underlined. Insertion sites of T-DNA alleles tash3-1, tash3-2 and tash3-3, as well as the site of CRISPR/Cas9-induced mutation in tash3-c, are indicated. (B) Seedlings of tash3-c and tash3-3 single mutants and in combinations with sh3p12310G. For clarity, the rightmost panel shows sh3p12310G tash3-3 quadruple homozygotes after the removal of lower order mutant seedlings from a population segregating for sh3p1. Graph shows main root lengths of individual seedlings after 9 d of in vitro growth from a representative experiment. Col-0: 5.80 ± 0.58 cm (mean ± s.d.), n=28; tash3-c: 6.08 ± 0.49 cm, n=24; tash3-3: 5.26 ± 0.51 cm, n=27; sh3p12310G: 3.16 ± 0.41 cm, n=26; sh3p12310G tash3-c: 3.33 ± 0.44 cm, n=27; sh3p12310G tash3-3: 2.24 ± 0.43 cm, n=17. Values were compared using One-way ANOVA (P<0.00001) with post-hoc Tukey HSD test, groups of significantly different values are indicated. (C) Adults of tash3-c and tash3-3 single mutants and in combinations with sh3p12310G. Inset shows a magnified view of sh3p12310G tash3-3. (D) TIRF images of CLC2-GFP in the epidermis of early elongation zone of seedling roots of wild type, sh3p12310G tash3-3, and lower order mutants in a population segregating for sh3p1. Scale bar – 2 μm. Graph shows a quantification of CLC2-GFP foci density, each data point represents one root. Wild type: 113.9 ± 21.0 foci per 100 μm2 (mean ± s.d.), n=30; SH3P1 and sh3p1 +/-sh3p23 -/-tash3-3 -/-: 88.4 ± 18.9 foci per 100 μm2, n=32; sh3p12310G tash3-3: 65.6 ± 19.8 foci per 100 μm2, n=30. Values were compared using One-way ANOVA (P<0.00001) with post-hoc Tukey HSD test, groups of significantly different values are indicated.

In contrast with *tash3-1* and *tash3-2*, both *tash3-3* and *tash3-c* were viable as homozygotes, but surprisingly, exhibited distinct phenotypes. While *tash3-c* was indistinguishable from the wild type during seedling and adult development (Figure 5B and 5C), *tash3-3* had slightly reduced seedling root lengths (Figure 5B) and clearly deficient adult development, with a reduced number of stems, reduced fertility, and early senescence of rosette leaves (Figure 5C). Since both alleles delete the SH3 domain, the defects exhibited by *tash3-3* indicate additional effects of the T-DNA insertion, presumably reduced gene expression. We tested *TASH3* expression levels in *tash3-3 and tash3-c* by qPCR, and found that both alleles had reduced transcript levels, but the expression in *tash3-3* was lower than in *tash3-c* across biological replicates and experiment repetitions (Suppl. Figure 2B). It might be that *TASH3* transcript levels explain the phenotypic difference between mutant alleles, but additional effects on translation from the abnormal *tash3-3* mRNA, presumably containing parts of the T-DNA sequence, or effects on tash3-3 protein stability, may contribute. Overall, the normal phenotype of *tash3-c* asserts that the SH3 domain of TASH3 has no unique, detectable function. By comparison, the *tash3-3* phenotype cannot be explained by the loss of the SH3 domain alone, and we interpret it as a partial loss-of-function of TASH3 as a whole, and by extension, as a partial loss-of-function of the TPLATE complex in which it is a core component.

We introduced both *tash3-3* and *tash3-c* into *sh3p123*^*10G*^ to assess genetic interactions. A potential redundant function of the SH3 domains of TASH3 and SH3P1-3 could be tested in *sh3p123*^*10G*^ *tash3-c*. This quadruple homozygote exhibited phenotypes identical to *sh3p123*^*10G*^ both at seedling and adult stages of development (Figure 5B, 5C), indicating that the SH3 domain of TASH3 does not contribute to a function redundant with SH3P1-3. In turn, *sh3p123*^*10G*^ *tash3-3* quadruple mutants had diminished seedling root growth rates in comparison to both parental lines (Figure 5B), while as adults, they exhibited a striking phenotype characterized by the development of very small plants with reduced rosette leaves, rarely bolting, and producing only up to a few flowers on short inflorescence stems (Figure 5C). These plants were infertile and the line was maintained as *sh3p1* heterozygote.

To assess rates of CME in *sh3p123*^*10G*^ *tash3-3*, we introduced the clathrin marker CLC2-GFP and performed TIRF microscopy in the early elongation zones of seedling roots. Segregating plants wild type or heterozygous for *sh3p1* had a mild decrease in CCP density compared with the wild type (Figure 5D). This phenotype is typical to *tash3-3*, as it was observed in an independent set of TIRF experiments with CLC2-mRuby expressed in *tash3-3*. single mutant (Adamowski et al., 2022b). In turn, *sh3p123*^*10G*^ *tash3-3* quadruple homozygotes had a strongly decreased density of CCPs at the PM (Figure 5D). This demonstrates a contribution of SH3P1-3 to gross endocytic activity, not detected when SH3P1-3 function alone was abolished (Figure 1D), but manifested in a sensitized genetic background.

Taken together, we did not find evidence for a common function of the SH3 domains of TASH3 and SH3P1-3, as shown by the lack of effects of the *tash3-c* allele, where the SH3 domain of TASH3 is abolished with a relative specificity. The chance isolation of *tash3-3* as a partial loss-of-function of the TPLATE complex revealed a role of SH3P1-3 in gross CME, masked in the original mutant, and manifested by strong deficiencies in overall growth and development, as well as in CCP formation, in the mutant cross.

## Discussion

### SH3P1-3 are endophilin/amphiphysin homologues engaged in a novel interaction with auxilin-likes

Here, we characterize the BAR-SH3 domain proteins SH3P1-3 of *A. thaliana* as homologues of endophilin and amphiphysin, which play important functions at late stages of CME in non-plant systems (Shupliakov et al., 1997; Milosevic et al., 2011). SH3P1-3 dynamically localize to discrete foci at the PM, and colocalization indicates specific recruitment at the late stage of CCP formation in a subset of CME events, similarly to non-plant counterparts (Taylor et al., 2011). Previously reported protein-protein interactions (Lam et al., 2001; Adamowski et al, 2018), the remarkable level of spatio-temporal colocalization at the PM, and the reaction of SH3P2-GFP to inhibition of CME by AUXILIN-LIKE1 overexpression, all indicate a functional association of SH3P1-3 with the putative auxilin-like uncoating factors AUXILIN-LIKE1/2. This interaction promotes AUXILIN-LIKE1/2 localization at the PM, likely including the late recruitment at CCPs, as seen by the loss of AUXILIN-LIKE1 from the PMs in *sh3p123* mutant. Yet, parallel mechanisms may contribute to AUXILIN-LIKE1/2 recruitment to CCPs, since events of such recruitment could be still detected in *sh3p123*, although a quantitative comparison with the wild type was not feasible. Nevertheless, taken as a whole, our findings provide strong support for a novel, plant-specific interaction module in CME, where BAR-SH3 domain-containing proteins engage with auxilin-likes, rather than with synaptojanins, most likely to promote vesicle uncoating.

Beside AUXILIN-LIKE1/2, three other auxilin-like proteins, AUXILIN-LIKE3-5, are proposed to contribute to the uncoating process in *A. thaliana*, where they are accompanied by ARK1-LIKE, a putative homologue of yeast protein kinases that mediate CCV uncoating through the phosphorylation of adaptors (Adamowski et al., 2022a). Of these, ARK1-LIKE may interact with SH3P1-3, as suggested by a yeast two-hybrid interaction between SH3P1 with ARK1-LIKE and the presence of a Pro-rich domain in ARK1-LIKE (Lam et al. 2001). Associations of SH3P1-3 with AUXILIN-LIKE3-5 are unlikely, as these auxilin homologues do not possess Pro-rich domains, while AUXILIN-LIKE3 does not localize to the PM, but may be involved in late uncoating of CCVs upon arrival at the early endosome, a recently described feature of plant CME (Narasimhan et al., 2020; Adamowski et al., 2022a).

### Other potential functions of SH3P1-3 in endocytosis

The mutant phenotype of *sh3p123*, compared with a lack of observable deficiencies in *auxilin-like1/2* or *auxilin-like1/2 ark1-like* (Adamowski et al., 2018, 2022a) indicates that the function of SH3P1-3 encompasses additional activities beside the recruitment of AUXILIN-LIKE1/2, and possibly ARK1-LIKE. Notable are the previously reported interactions of SH3P1-3 with dynamins, analogical to the mechanism of action of endophilin and amphiphysin in non-plant models. SH3P1 and SH3P2 were reported to interact with DYNAMIN RELATED PROTEIN 1A (DRP1A) (Ahn et al., 2017; Baquero Forero and Cvrčková, 2019), possibly indirectly, given that DRP1A does not possess Pro-rich regions. In turn, SH3P3 interacts with DRP2A, which, like its homologue DRP2B, contains Pro-rich domains (Lam et al., 2002). If SH3P1-3 are required for the activity of DRP2, this contribution may be only partial, as otherwise, *sh3p123* would be expected to manifest gametophytic lethality like *drp2ab* (Backues et al., 2010). Beside this, it may be speculated that SH3P1-3 participate in a clathrin-independent mode of endocytosis, like endophilin, which acts in Fast Endophilin-Mediated Endocytosis (Casamento and Boucrot, 2020). Localization of SH3P2-GFP to PM sites not containing clathrin is supportive of this possibility, but the likely presence of AUXILIN-LIKE1 there, and the apparent sensitivity of all PM localization of SH3P2-GFP to clathrin silencing, argue against this scenario. Finally, knock-out mutants isolated in this study call for a verification of functions assigned to SH3P2 in part through the lethal phenotypes of RNAi lines silencing *SH3P2* (Zhuang et al., 2013; Ahn et al., 2017).

### Implications of the genetic interaction with the TPLATE complex

Quantitatively, the function of SH3P1-3 in CME was best revealed in a genetic interaction between *sh3p123* and *tash3-3*. We interpret *tash3-3* as a partial loss-of-function of the TPLATE complex, rather than specifically affecting the SH3 domain of its subunit TASH3, through comparison with *tash3-c* where knock-out of this SH3 domain did not lead to any evident deleterious effects. The strong deficiencies in growth and CME in *sh3p123 tash3-3* demonstrate a contribution of SH3P1-3 to CME, but these observations also lead to questions of a general nature, regarding the possible courses that CCP formation can take and function effectively. Genetic analysis indicates that CME can function well without SH3P1-3, but only if TPLATE complex is fully active *(sh3p123)*, or with only partial TPLATE complex function, provided that SH3P1-3 are present instead (*tash3-3)*. In which sense are SH3P1-3 redundant with the TPLATE complex, if the TPLATE complex acts in CME from the early stages, likely as an adaptor (Gadeyne et al., 2014), while SH3P1-3 are recruited to CCPs late, contributing to uncoating, and potentially, scission? Currently, the only commonality between the two that may explain this redundancy, are physical interactions with dynamins (Lam et al., 2002; Gadeyne et al., 2014; Ahn et al., 2017; Baquero Forero and Cvrčková, 2019). On the other hand, if the TPLATE complex and SH3P1-3 contribute to this redundant function by distinct molecular activities, a situation emerges where CME may proceed effectively through different molecular mechanisms of vesicle formation, provided by coats of distinct compositions. This and similar examples of genetic interactions in CME (Adamowski et al., 2022b) suggest that such flexibility of the endocytic process is a likely scenario.

## Materials and methods

### Plant material

The following previously described *A. thaliana* lines were used in this study: *CLC2*_*pro*_*:CLC2-GFP* (Konopka et al., 2018), *UBQ10*_*pro*_*:mCherry-AUXILIN-LIKE1, CLC2*_*pro*_*:CLC2-GFP UBQ10*_*pro*_*:mCherry-AUXILIN-LIKE1, XVE»AUXILIN-LIKE1* (Adamowski et al., 2018), *XVE»amiCHCa* (Adamowski and Friml, 2021), *UBQ10*_*pro*_*:SH3P2-GFP* (Zhuang et al., 2013), *sh3p3* (SALK_065970; Nagel et al., 2017). Lines generated as part of this study with details of mutant alleles are listed in Suppl. Table 1 and primers used for genotyping in Suppl. Table 2.

### *In vitro* cultures of *A. thaliana* seedlings

Seedlings were grown in *in vitro* cultures on half-strength Murashige and Skoog (1/2MS) medium of pH=5.9 supplemented with 1% (w/v) sucrose and 0.8% (w/v) phytoagar at 21°C in 16h light/8h dark cycles with Philips GreenPower LED as light source, using deep red (660nm)/far red (720nm)/blue (455nm) combination, with a photon density of about 140μmol/(m^2^s) +/-20%. β-estradiol (Sigma-Aldrich) was solubilized in 100% ethanol to 5 mg/mL stock concentration and added to ½MS media during preparation of solid media to a final concentration of 2.5 μg/mL. Seedlings of *XVE»AUXILIN-LIKE1* and *XVE»amiCHCa* lines were induced by transferring to β-estradiol-supplemented media at day 3. Petri dishes for TIRF imaging in hypocotyls of etiolated seedlings were initially exposed to light for several hours and then wrapped in aluminium foil.

### Confocal Laser Scanning Microscopy

4 to 5 d old seedlings were used for live imaging with Zeiss LSM800 confocal laser scanning microscope with 20X lens. Measurements of relative PM signal intensities of SH3P2-GFP and mCherry-AUXILIN-LIKE1 were performed in Fiji (https://imagej.net/Fiji) as a ratio between mean grey value of a line drawn over multiple PMs in each CLSM image, and a rectangle covering the whole RAM visible. The *UBQ10*_*pro*_*:mCherry-AUXILIN-LIKE1* transgene exhibited silencing, which was particularly visible in the *sh3p123* cross; the experiments were thus conducted on seedlings pre-selected under a stereomicroscope equipped with a UV lamp as expressing relatively brighter mCherry signals. Images in both genotypes were captured by CLSM with identical detection settings to ascertain validity of the comparison.

### Total Internal Reflection Fluorescence microscopy

Early elongation zone of roots in excised ∼1 cm long root tip fragments from 7d old seedlings, as well as apical ends of excised hypocotyls from 3d old etiolated seedlings, were used for TIRF imaging. Imaging was performed with Olympus IX83 TIRF microscope, using a 100X TIRF lens with an additional 1.6X or 2X magnification lens in the optical path. Time lapses of 100 frames at 0.5 s or 1 s intervals with exposure times of 195 ms or 200 ms, or single snapshots of 200 ms exposure, were taken, depending on the experiment. Two-channel time lapses were captured sequentially. CLC2-GFP foci were counted in square regions of 36 μm^2^ taken from the captured TIRF images or movies using Fiji (https://imagej.net/Fiji).

### Molecular cloning

All constructs generated in this study are listed in Suppl. Table 3 and primers used for cloning in Suppl. Table 2. Sequences of *SH3P1, SH3P3*, and *CLC2* were cloned into pENTR/D-TOPO and pDONR221 entry vectors (Invitrogen). *35S:SH3P1-GFP* and *35S:SH3P3-GFP* were generated in pH7FWG2 expression vectors (Karimi et al., 2002) by LR Clonase II and used for transformation of *sh3p123* alongside the previously cloned 35S:SH3P2-GFP/pH7FWG2 (Adamowski et al., 2018). *RPS5A:CLC2-mRuby* was generated in pK7m34GW expression vector (Karimi et al., 2002) by LR Clonase II Plus by combining RPS5A/pDONRP4P1r, CLC2/pDONR221, and mRuby3/pDONRP4P1r.

### CRISPR/Cas9 mutagenesis

CRISPR mutagenesis of *SH3P1, SH3P2*, and *TASH3* was performed with the use of pHEE401 binary vector and template plasmids pCBC-DT1T2 and pCBC-DT2T3 (Wang et al., 2015). sgRNA sequences were selected with the use of CRISPR RGEN Tools website (http://www.rgenome.net/cas-designer/). sgRNA sequences used in each construct are given in Suppl. Table 3. Mutagenesis of *SH3P1* and *SH3P2* was performed in Col-0 background using SH3P-A CRISPR/pHEE401 and in *sh3p3* background using SH3P-B CRISPR/pHEE401. Mutagenesis of TASH3 was performed in Col-0 and in *sh3p123*^*10G*^ backgrounds independently but identical mutant allele *tash3-c* was isolated. In T1 plants, target sequences were PCR-amplified and sequenced using primers listed in Suppl. Table 2. Where homozygotes were not found, further genotyping was performed in T2 generation. Plants negative for the CRISPR/Cas9 transgene were selected in T2 generation by PCR on *Cas9* gene sequence. Details of generated mutant alleles are given in Suppl. Table 1.

### Quantitative reverse transcriptase PCR

Total RNA was isolated from 5d old seedlings using RNeasy Plant Mini Kit (Qiagen). cDNA was synthetised using iScript™ cDNA Synthesis Kit (Bio-Rad). qPCR was performed with Luna^®^ reagent (New England Biolabs) in Roche LightCycler® 480. *TUB2* and *PP2AA3* were used as reference genes. Primer sequences are listed in Suppl. Table 2.

### Accession numbers

Sequence data from this article can be found in the GenBank/EMBL libraries under the following accession numbers: SH3P1 (AT1G31440), SH3P2 (AT4G34660), SH3P3 (AT4G18060), TASH3 (AT2G07360), AUXILIN-LIKE1 (AT4G12780), CLC2 (AT2G40060).

## Author Contributions

M.A. and J.F. designed research, analysed data and wrote the manuscript. M.A. and I.M. performed research. M.N. generated *RPS5A:CLC2-mRuby*.

## Acknowledgements

The authors wish to acknowledge Dr Daniel van Damme for mRuby3/pDONRP2rP3 and Prof. Qi-Jun Chen for sharing plasmids used for CRISPR/Cas9 mutagenesis. This work was supported by the Austrian Science Fund (FWF): I 3630-B25.

---

**Suppl. Figure 1.**
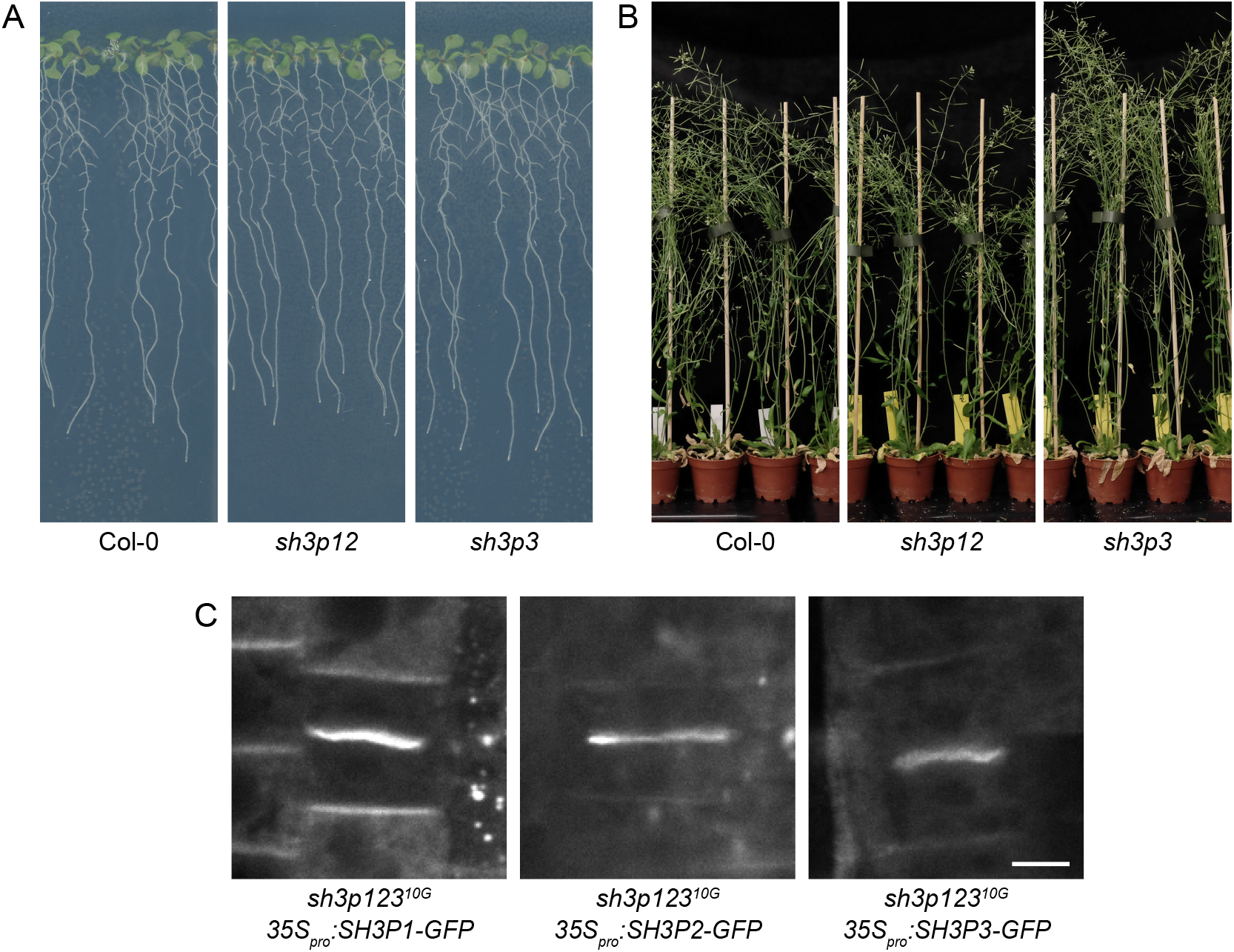
Additional data on sh3p mutants and SH3P localization (A) Normal seedling development of sh3p12 and sh3p3 mutants. (B) Normal adult development of sh3p12 and sh3p3 mutants. (C) CLSM images of SH3P1-GFP, SH3P2-GFP, and SH3P3-GFP in seedling RAMs of complemented sh3p12310G mutants showing strong binding to cell plates. Scale bar – 5 μm.

**Suppl. Figure 2.**
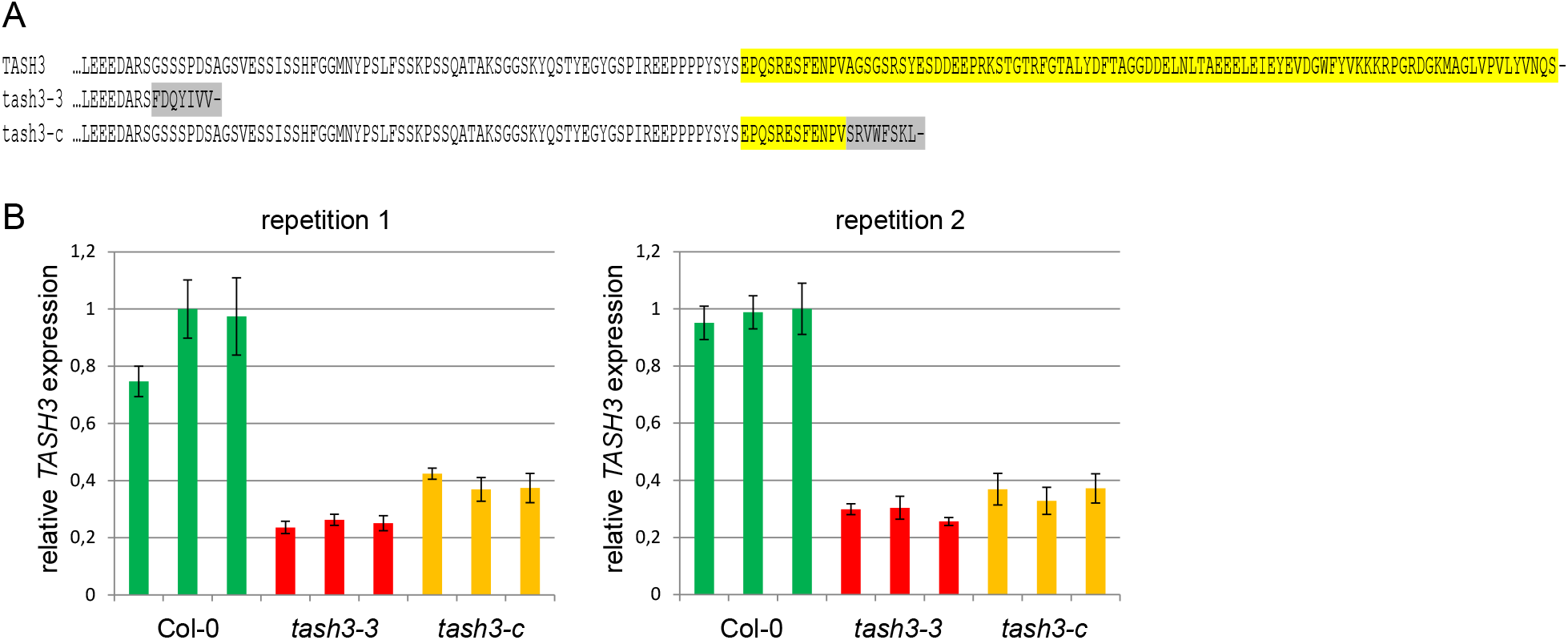
Additional data on tash3 mutants (A) Protein sequence alignment of the C-termini of TASH3 and predicted mutant proteins tash3-3 and tash3-c. tash3-3 sequence is predicted using a sequenced tash3-3 T-DNA insertion site. Sequence marked in yellow encodes the SH3 domain. Sequences in grey are the altered terminal sequences of tash3-3 and tash3-c. (B) Two repetitions of a qPCR experiment assessing the expression of TASH3 in tash3-3 and tash3-c. TASH3 primer pair recognizes an mRNA region shortly upstream of the T-DNA insertion in tash3-3. Each experiment involved three biological replicates of each genotype, shown as three values on the graphs. Error bars indicate standard deviations of three technical replicates. TUB2 and PP2AA3 were used as reference genes.

