## Supplemental Tables for "SH3Ps recruit auxilin-like vesicle uncoating factors into clathrin-mediated endocytosis"

Suppl. Table 1. Lines generated in this study.

| **Line name** | **Notes** |
| --- | --- |
| *sh3p1,2* | *SH3P1* …GCTGCTAAGgta…  *sh3p1* …GCTGACTAAGgta…  *SH3P2* …ACAAGCTAGCAG…  *sh3p2* …ACAAGCTTAGCAG… |
| *sh3p3* | SALK_065790 |
| *sh3p1,2,3^9C (9C-Δ14)^* | *SH3P1* …agCGTTTGCAAAGGAAC…  *sh3p1* …agCGAAGAGGAAC…  *SH3P2* …AGTTGCGAGACAGCAGCA…  *sh3p2* …AGCA… |
| *sh3p1,2,3^10G (10G-4E)^* | *SH3P1* …agCGTTTGCAAAGGAAC…  *sh3p1* …agCGTTTGGCAAAGGAAC…  *SH3P2* …GAGACAGCAGCAGgtta…  *sh3p2* …GAGACAGCAAGCAGgtta… |
| *sh3p1,2,3^10G^ CLC2_pro_:CLC2-GFP UBQ10_pro_:mCherry-AUXILIN-LIKE1* |  |
| *tash3-1* +/- | SALK_122269 |
| *tash3-2* +/- | SALK_123877 |
| *tash3-3* | SALK_011079 |
| *sh3p1*+/-*,2,3^10G^ tash3-3* |  |
| *sh3p1*+/-*,2,3^10G^ tash3-3 CLC2_pro_:CLC2-GFP*  (silenced *UBQ10_pro_:mCh-AUXILIN-LIKE1*) |  |
| *tash3-c* | *TASH3* …CCTGTAGCAGG…  *tash3-c* …CCTGTAAGCAGG… |
| *sh3p1,2,3^10G^ tash3-c* | *TASH3* …CCTGTAGCAGG…  *tash3-c* …CCTGTAAGCAGG… |
| *RPS5A_pro_:CLC2-mRuby* | Kan |
| *UBQ10_pro_:SH3P2-GFP RPS5A_pro_:CLC2-mRuby* | Kan |
| *UBQ10_pro_:SH3P2-GFP UBQ10_pro_:mCherry-AUXILIN-LIKE1* | Kan, Basta |
| *35S_pro_:SH3P1-GFP* | Hyg |
| *35S_pro_:SH3P3-GFP* | Hyg |
| *sh3p1,2,3^10G^ 35S_pro_:SH3P1-GFP* | Hyg |
| *sh3p1,2,3^10G^ 35S_pro_:SH3P2-GFP* | Hyg |
| *sh3p1,2,3^10G^ 35S_pro_:SH3P3-GFP* | Hyg |
| *UBQ10_pro_:SH3P2-GFP XVE»AUXILIN-LIKE1* |  |
| *UBQ10_pro_:SH3P2-GFP CAP1_pro_:CAP1-mCh XVE»amiCHCa* |  |

Suppl. Table 2. Primers used in this study

| **Primer name** | **Sequence** | **Purpose** |
| --- | --- | --- |
| SH3P1-TOPO-F | CACCATGGAAGCTATAAGAAAGCAAGC | cloning |
| SH3P1-TOPO-R | TCACTGTTGCTTGGAGTTTGA | cloning |
| attB1-SH3P1-F | GGGGACAAGATTGTACAAAAAAGCAGGCTATGGAAGCTATAAGAAAGCAAGC | cloning |
| attB2-SH3P1-Rns | GGGGACCACTTTGTACAAGAAAGCTGGGTACTGTTGCTTGGAGTTTGATTC | cloning |
| SH3P3-TOPO-F | CACCATGGATGCGTTTAGAAGACAA | cloning |
| SH3P3-TOPO-R | TCAGTAAACTTCAGCAGCAAAG | cloning |
| SH3P3-TOPO-Rns | GTAAACTTCAGCAGCAAAGTTG | cloning |
| sh3p3-F | GGCTCAGACACATTGAAGCA | *sh3p3* genotyping |
| sh3p3-R | acgcggtaagaccaaaagtg |  |
| LBb1.3 | ATTTTGCCGATTTCGGAAC | SALK T-DNA lines genotyping |
| SH3P1-cF | ttgggggtagataggattgttg | *sh3p1* CAPS genotyping, differential digestion with HpyCH4V in all alleles |
| SH3P1-cR | AGAGTTTCCCTCTCGCCTTC |  |
| SH3P2-cF | tcgtgttcgaaagcagagaa | *sh3p2* CAPS genotyping, differential digestion with Fnu4HI in all alleles |
| SH3P2-cR | ccctttgacaaaaacttctcca |  |
| tash3-1-F | acgtgcatctcacgtgctt | *tash3-1* genotyping |
| tash3-1-R | GGCCAGAGAAGGATCAACTG |  |
| tash3-2-F | ggcccgagttaaatgttatttg | *tash3-2* genotyping |
| tash3-2-R | ggcagcagaaagaggttgat |  |
| tash3-3-F | atggggtcaagaatgaatgc | *tash3-3* genotyping |
| tash3-3-R | AGCGCTGTTCCAAATCTTGT |  |
| tash3-cF | AAAGCCTTCCTCACAAGCAA | CRISPR genotyping (sequencing) |
| tash3-cR | tgatcccggttagtttacgg |  |
| TASH3-qF | GGTTGGGTATGATGACATGTGG | qPCR |
| TASH3-qR | CGCATCTTCTTCCTCCAATTCT | qPCR |
| TUB2-qF | AAACTCACTACCCCCAGCTTTG | qPCR |
| TUB2-qR | CACCAGACATAGTAGCAGAAATCAAGT | qPCR |
| PP2AA3-qF | TAACGTGGCCAAAATGATGC | qPCR |
| PP2AA3-qR | GTTCTCCACAACCGCTTGGT | qPCR |
| CAS9-F | cggcctcgatattgggactaactct | CRISPR/Cas9 T-DNA detection |
| CAS9-R | cttatctgtggagtccacgagcttc | CRISPR/Cas9 T-DNA detection |

Suppl. Table 3. Plasmids generated in this study.

| **Plasmid name** | **Notes** |
| --- | --- |
| SH3P1/pENTR/D-TOPO | with STOP codon |
| SH3P1/pDONR221 | without STOP codon |
| SH3P3/pENTR/D-TOPO | with STOP codon |
| SH3P3/pENTR/D-TOPO | without STOP codon |
| CLC2/pDONR221 | without STOP codon |
| 35S:SH3P1-GFP/pH7FWG2 |  |
| 35S:SH3P3-GFP/pH7FWG2 |  |
| SH3P-A CRISPR/pHEE401 | Used for mutagenesis in Col-0 background  SH3P1-A AGCTCTACTAAAGCTGCTA  SH3P2-A AATTAGAAAACAAGCTAGC |
| SH3P-B CRISPR/pHEE401 | Used for mutagenesis in *sh3p3* background  SH3P1-B ctgttcatagCGTTTGCAA  SH3P2-B CAAGTTGCGAGACAGCAGC |
| TASH3 CRISPR/pHEE401 | 1 CAAACGATTCACGACTTTG  2 GAGAACCAGACCCTGCTAC  3 TCTATGACTTCACAGCAGG |
| RPS5A:CLC2-mRuby/pK7m34GW |  |
